# Mapping DNA replication with nanopore sequencing

**DOI:** 10.1101/426858

**Authors:** Magali Hennion, Jean-Michel Arbona, Corinne Cruaud, Florence Proux, Benoît Le Tallec, Elizaveta Novikova, Stefan Engelen, Arnaud Lemainque, Benjamin Audit, Olivier Hyrien

## Abstract

We have harnessed nanopore sequencing to study DNA replication genome-wide at the single-molecule level. Using *in vitro* prepared DNA substrates, we characterized the effect of bromodeoxyuridine (BrdU) substitution for thymidine on the MinION nanopore electrical signal. Using a neural-network basecaller trained on yeast DNA containing either BrdU or thymidine, we identified BrdU-labelled tracts in yeast cells synchronously entering S phase in the presence of hydroxyurea and BrdU. As expected, the BrdU-labelled tracts coincided with previously identified early-firing, but not late-firing, replication origins. These results open the way to high-throughput, high-resolution, single-molecule analysis of DNA replication in many experimental systems.

## INTRODUCTION

DNA replication is the biological process by which a genome is accurately duplicated before a cell divides into two daughter cells. Eukaryotic organisms replicate their genome from multiple, stochastically activated replication origins. Origin activation establishes bidirectional replication forks that progress along and duplicate the DNA until they merge with forks emanating from adjacent origins. Understanding the regulation of these events is essential as their perturbations threaten genome stability.

In the last decade, DNA microarray and massive DNA sequencing techniques have triggered an explosion of genome-wide replication mapping studies. However, these cell-population based methods provide an average profile of DNA replication which masks cell-to-cell heterogeneity in origin activation and fork progression. Single-molecule methods can reveal this heterogeneity. Typically, cells are consecutively pulsed with two thymidine analogues that are incorporated into newly synthesized DNA; total genomic DNA is then extracted, spread on microscope glass coverslips, and the labelled tracts are detected with appropriate fluorescent antibodies [1, 2]. This approach allows to visualize the progression of individual replication forks during the pulses and to infer the spacing of initiation and termination events. However it does not provide DNA sequence information unless combined with fluorescent *in situ* hybridization with specific DNA probes [3]. This is technically difficult and limits the analysis to a tiny portion of the genome. Recently, significant improvements in throughput and automation of single-molecule analysis were achieved by (i) using fluorescent dNTPs to directly label the newly replicated DNA; (ii) barcoding total DNA by fluorescent labeling at nicking endonuclease cutting sites and (iii) stretching labelled DNA in nanochannel arrays originally developed for automated genome assembly (Bionano Genomics, [4, 5]). Nevertheless, nanochannel image analysis and genomic alignment of barcoded DNA requires complex bioinformatic processing and the resolution of these methods is still limited to ~1 kb due to optical microscopy limitations and DNA stretching inhomogeneities.

A novel sequencing technology, namely nanopore sequencing, has the potential to bypass these limitations. In MinION nanopore sequencing (Oxford Nanopore Technologies, ONT), a long single strand of DNA (up to 2.3 Mb [6]) is translocated by a molecular motor through a protein nanopore inserted in a voltage-biased membrane separating two ionic solution-filled chambers. The ionic conductivity through the nanopore is particularly sensitive to the nucleobases located in its narrowest region. Thus, changes in the ionic current levels during translocation reveal changes in the DNA sequence. Several consecutive nucleobases in the narrowest region can influence the ionic current. Translating a sequence of current values into a DNA sequence is therefore a non-trivial task. Several machine learning approaches using either Hidden Markov Models (HMM) [7, 8] or Recurrent Neural Networks (RNN) [9, 10] have been developed by ONT or by academic groups to reach this goal. Importantly, such approaches can discriminate minor bases modifications such as cytosine methylation and hydroxymethylation [11, 12]. These results suggest that nanopore sequencing may allow direct detection of modified nucleobases incorporated in newly replicated DNA. This would simultaneously provide the sequence and replicative status of long, native DNA fragments at near-nucleotide resolution.

Here, we demonstrate that this approach is feasible. We can directly detect bromodeoxyuridine (BrdU), a thymidine analogue widely used in DNA replication studies, both in synthetic test templates and following *in vivo* replicative incorporation in the yeast *Saccharomyces cerevisiae*. Replicative stretches synthesized at the onset of S phase and detected in this manner map to origins known to be active in early S phase but not to later-activated origins. This provides, to the best of our knowledge, the first proof-of-principle that nanopore sequencing can be used to map replication genome-wide at the single-molecule level, surpassing the throughput of alternative optical methods and promising DNA replication analysis with unprecedented resolution.

## RESULTS

### *In vitro* templates

In order to measure the effect of BrdU incorporation on the nanopore electric signal, we first generated control or BrdU-hemisubstituted DNA duplexes using a single primer extension of linearised plasmid DNA in the presence of either dTTP or BrdUTP, followed by exonuclease degradation of the non-template strand (Fig. 1a). Bioanalyzer electrophoretic analysis and Qubit quantification of the purified reaction products (Fig. 1b) revealed a high yield of primer extension and an electrophoretic shift with respect to the starting duplex plasmid associated with BrdUTP but not dTTP incorporation. The small amount of duplex DNA observed in the absence of dTTP and BrdUTP likely resulted from partial renaturation of the template before exonuclease degradation. The primer extension products were sequenced using the ONT MinION (R9 chemistry) and the "2D" protocol where the two strands of a DNA duplex are consecutively read thanks to a hairpin adapter. We obtained 115K and 77K reads for the dTTP and the BrdUTP sample, respectively (sequencing information is summarized in Table S1).

The raw data were basecalled using Metrichor (ONT) and the resulting sequences were aligned to the plasmid sequence using BWA MEM [13] with parameters adapted to the error rate (see Materials and Methods, Supp. Fig. S1) and considering the two complementary DNA strands independently. As Metrichor was devised to detect canonical bases, the presence of BrdU could potentially affect basecalling and subsequent mapping. Indeed, the percentage of mapped reads for the BrdUTP sample was lower (49%) than for the dTTP sample (60%; Table S2). Metrichor classifies the reads into "pass" and "fail" categories based on the presence of a second strand read and its similarity to the first strand (2D protocol). The fraction of 2D reads was similar for both samples (57% vs. 59%) but the percentage of "pass" reads was lower for BrdUTP (13%) than for dTTP (18%), indicating lower complementarity of the two strand reads. Importantly, 99% of the "pass" reads were mapped for both samples (Supp. Fig. S1b and Table S2). However, looking at strand-oriented miscalls within the "pass" mapped reads, the parental (R) and newly replicated (F) strand miscall rates were similar for the dTTP sample (7% vs. 7%) but different for the BrdU sample (7% vs. 11%, respectively, Supp. Fig. S1c), confirming that BrdU affects the current in the pore. Together, these results suggest that current alterations due to BrdU reduce Metrichor basecalling accuracy but to an extent that does not strongly affect alignment of BrdU-substituted reads to the reference sequence. Python scripts were then developed to realign the measured current intensity to the plasmid DNA sequence. To allow quantitative comparisons between multiple experiments, we first normalized each profile by subtracting its mean current intensity. As exemplified on Fig. 1 (c and d), the presence of BrdU increased the current intensity at many, though not all, T positions. Since T was generally associated with the strongest current values in native DNA, this may explain why the current perturbation associated with BrdU did not result in systematic base calling errors at T sites (Supp. Fig. S1c). Notably, the incorporated BrdU did not visibly perturb the current at other bases. We extracted for each pentamer the difference in median current value between BrdUTP and dTTP samples (Supp. Fig. S1d). Almost all forward-strand pentamers with a T in their middle displayed a positive current shift in the presence of BrdU, with 60% showing a shift > +3 pA. On the contrary, most pentamers lacking T exhibited a small negative shift in the BrdU sample, because a higher mean current than in the dTTP sample was subtracted during normalisation. Importantly, only the signal from the BrdU-substituted (F) strand was perturbed, while the complementary native (R) strand gave an identical signal to the thymidine control sample (Fig. 1d, Supp. Fig. S1d). A principal component analysis of current sequences corresponding to 1 kb showed a clear separation of BrdU-substituted and control fragments (Fig. 1e). This indicates that the absence of a strong BrdU-induced current shift at some positions does not preclude the detection of BrdU-labelled replicative tracts of hundreds of bases or more. The small number of reads from the BrdUTP sample that clustered with control reads were most likely true thymidine reads coming from a residual amount of native plasmid in the sample. Indeed, our parental strand degradation protocol did not seem to completely remove the native plasmid, with as much as 18% preserved in the absence of primer extension, most likely owing to strand reannealing (Fig. 1b). Overall, these results demonstrate that the presence of BrdU detectably alters the nanopore electric signal, with a signature that should become identifiable using appropriate machine learning.

**Figure 1:**
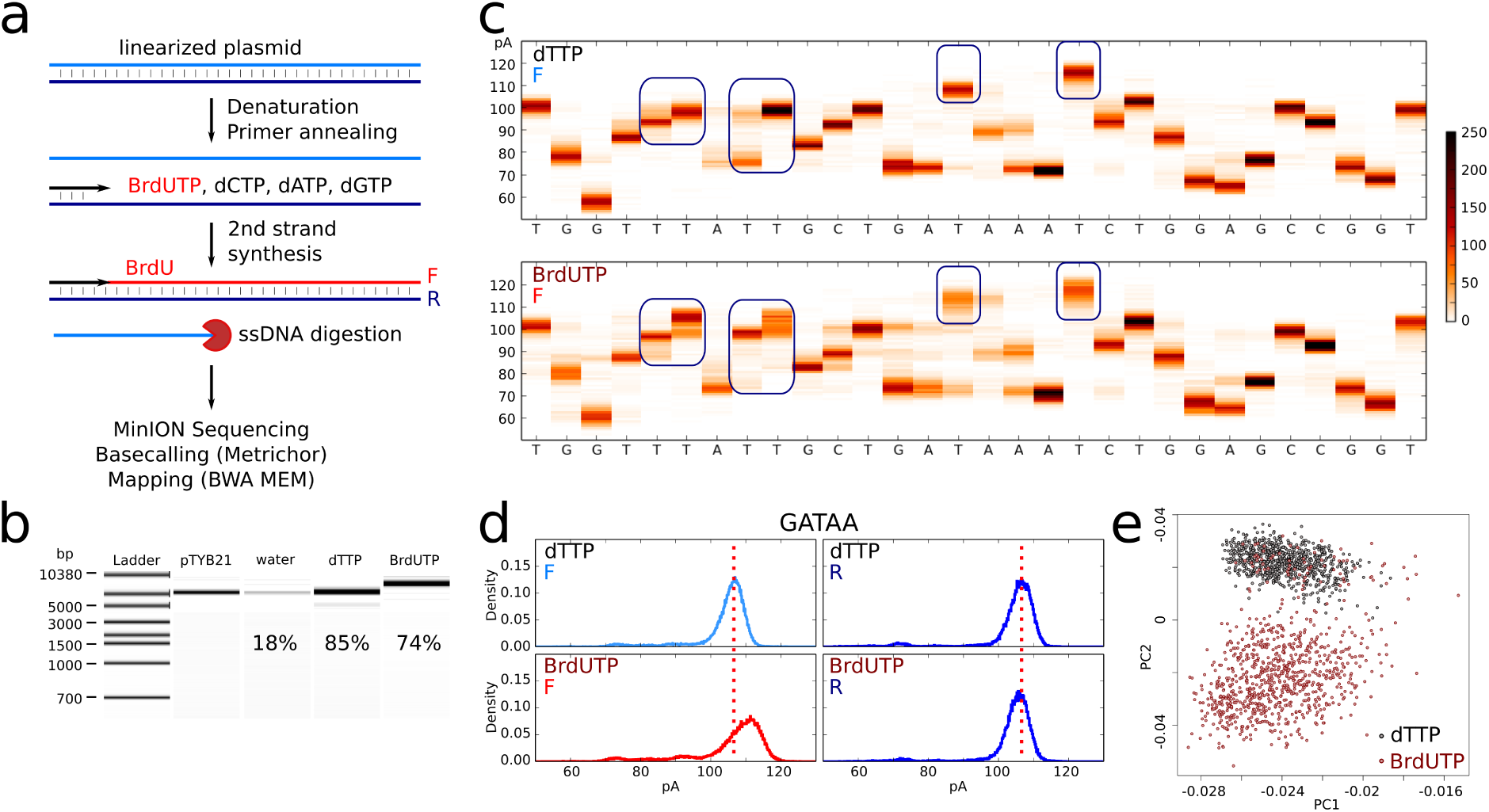
Effect of BrdU incorporation into DNA on nanopore sequencing current signal. a. Scheme of sample preparation. F, forward strand; R, reverse strand. b. Bioanalyzer size control of the samples, with Qubit yield indicated. pTYB21, linearised plasmid; water, primer extension in the absence of dTTP and Br-dUTP; dTTP, primer extension using canonical dNTPs; BrdUTP, primer extension using BrdUTP instead of dTTP. c. Example of a 30 bp sequence of the forward (F) strand (positions 1000-1029) with current distribution of 500 reads at each position. Upper panel: sample obtained using canonical dNTPs. Lower panel : dTTP was replaced by BrdUTP. Blue rectangles highlight some current shifts due to the presence of BrdU. BrdU did not induce a current shift at all thymidine sites. d. Current distribution for the ‘GATAA’ pentamer for the dTTP (top) and the BrdUTP (bottom) sample on the forward (F, modified strand, left) and the reverse (R, native strand, right) strands. e. Principal component analysis using current sequences from 1 kb read fragments from the dTTP (black) and BrdUTP (brown) samples (F strand). The first two components are represented. Only "pass" reads were used in c,d,e.

### ***In vivo*** BrdU incorporation

Metrichor was designed to call the 4 canonical DNA bases. Having shown that BrdU modified the nanopore signal on test templates prepared in *vitro*, we moved to an in *vivo* model to develop our own basecaller to detect BrdU as a fifth DNA base. A limiting step in machine learning is the quality of the training dataset. To distinguish thymidine from BrdU, one needs to train a model with a large diversity of DNA sequences containing only thymidine or only BrdU. To do so, we took advantage of the MCM869 yeast strain, which has been genetically modified to depend on exogenous thymidine (or a thymidine analogue) to replicate its genome [14]. We first verified that adding BrdU to the culture medium restored cell growth like thymidine (Supp. Fig. S2a), albeit at a slower rate (see Discussion). The cells stopped proliferating after 2-3 cycles, therefore we expected ~75% of genomic DNA strands to be totally substituted by BrdU, the remaining ~25% corresponding to the starting parental strands (Fig. 2a). We sequenced this BrdU-rich sample twice on the MinION (R9.4, 2D protocol), together with a control sample obtained from yeasts grown in the presence of thymidine. We obtained 20K-40K reads per sample (Table S3). The average read size was similar for all samples (Supp. Fig. S5a), indicating that BrdU incorporation did not substantially fragilize the DNA nor interfere with the sequencing. After basecalling by Metrichor the reads were aligned on the S288C genome by BWA MEM considering the two complementary strands separately. 79% and 82% of the thymidine and BrdU sample reads, respectively, were mapped. The improved mapping with respect to the plasmid experiment is likely attributable to the improved R9.4 chemistry. Current intensities were then realigned on the reference genome and, as seen for the *in vitro* template, the current was again positively shifted by the presence of BrdU (Fig. 2b).

We implemented RepNano, a recurrent neural network with an architecture similar to DeepNano [9], to convert the raw current from nanopore experiments into a DNA sequence (Supp. Fig. S3). RepNano allows the calling of 5 bases (A, T, G, C, B for BrdU) instead of the 4 canonical ones. To train the network, we used the Metrichor basecalls corrected for mistakes after alignment on the reference genome. RepNano learning was performed using a connectionist temporal classification objective function [15] that uses probabilistic rules insensitive to local misalignments between current plateaus and bases in the training sets. We trained a first RepNano model using 660 reads from the thymidine sample and 340 reads from the BrdU sample, assuming that all Ts were replaced by Bs (Model 1, Fig. 2c). This model was then used to basecall all the reads from the two samples in the 5 bases alphabet (Supp. Fig. S4a, Table S4). A false positive rate of 3.4% of B on T sites was observed in the thymidine reads while 69% of T sites were called B in the BrdU-enriched sample reads, in good agreement with our 75% expectation (Fig. 2a, Table S4). Looking at the percentage of B per read, we obtained a clear bimodal distribution allowing to classify most of the reads from the BrdU sample as either not substituted or fully substituted (Fig. 2d, Supp. Fig. S4b, Model 1). The RepNano-called reads were then mapped on the yeast genome after conversion of the Bs into Ts. The mappability of such called reads was slightly lower than with Metrichor (Supp. Fig. S5b, Table S5). To improve the model, we repeated the training with the same training dataset where the unsubstituted reads from the BrdU sample (BrdU content < 33%) were attributed to the thymidine sample (Fig. 2c, Supp. Fig. S4b, Model 2). False positive rate of the resulting RepNano Model 2 was reduced by a factor of 8 (0.4% of Bs in the thymidine sample) while B content slightly increased in the BrdU-rich sample (71%), compared to Model 1 (Table S4). Moreover we obtained fewer reads with intermediate B amount (Fig. 2d, Supp. Fig. S4b) and the mappability increased (Supp. Fig. S5b). We then represented all reads from a 250 kb genomic locus using a blue/red color map to show T/B density (Fig. 2e,f). The reads from the thymidine sample were completely blue, whereas most reads from the BrdU-rich sample were either fully blue (parental DNA) or fully red (BrdU-substituted DNA), as expected.

**Figure 2:**
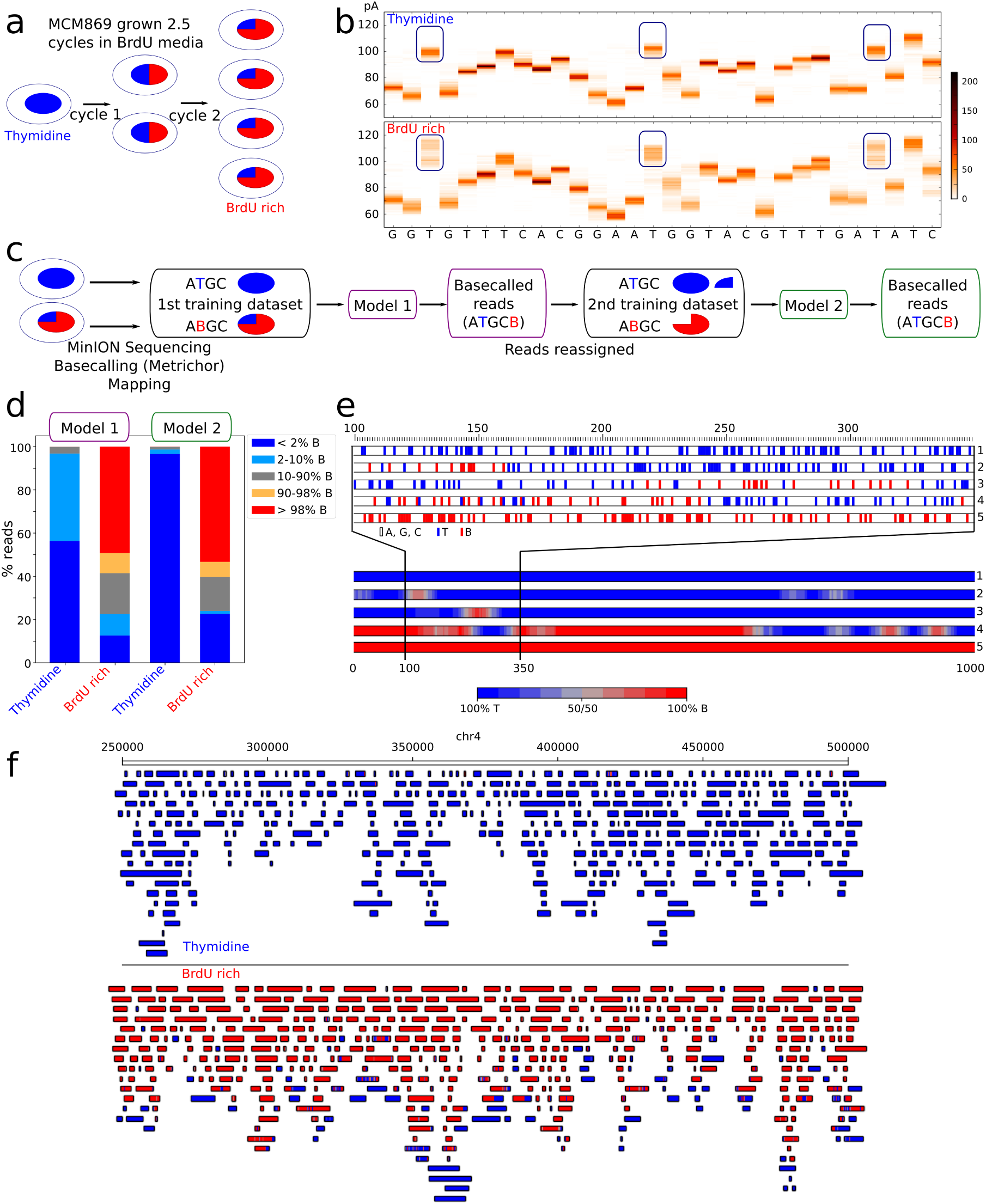
Detection of BrdU in yeast DNA. a. Scheme of the experiment: MCM869 yeasts were grown in BrdU medium for 2.5 cycles (according to OD measurements) resulting in a sample enriched in BrdU-DNA (red) but still containing native DNA (blue). b. Example of a 30 bp sequence (chr12:458000-458029, in rDNA repeats) with the current distribution at each position (about 400 reads). Blue rectangles highlight some current shifts due to the presence of BrdU. c. RepNano training and analysis pipeline. d. Comparison between the two models. The proportion of the reads with different amounts of B is represented for the thymidine and the BrdU enriched samples. e. Exemplary reads from the BrdU-enriched sample. Reads are represented as a heatmap of T vs. B density computed in 10 B/T sliding windows. f. Example of a 250 kb chromosomal segment with reads from the thymidine (top) and the BrdU rich (bottom) samples. Reads were vertically ordered according to their B density. Only "pass" reads were used in b,d,f.

### Early S phase labelling

*S. cerevisiae* replication origins are precisely positioned at sequences called ARSs (Autonomously Replicative Sequences) [16] and have been extensively studied. To validate our BrdU tract detection and mapping procedure we labelled early-firing origins by synchronizing cells in G1 and releasing them into S phase in the presence of BrdU and hydroxyurea (HU) (Fig. 3a). HU slows down replication forks and triggers a checkpoint that prevents late origin firing [17, 18, 19]. After 30 or 60 min, when cells were still in early S phase as seen by flow cytometry (Fig. 3b), a thymidine chase in the absence of HU was performed to allow the conversion of replication intermediates into dsDNA (Fig. 3a and Supp. Fig. S6). Similar protocols have been previously used to label early origins prior to microarray or sequencing analysis [20, 21, 22, 23, 24]. Genomic DNA was then extracted and sequenced on the MinION (R9.4), giving 416K and 44K reads for the 30 min and 60 min sample, respectively (Table S3). The reads were basecalled (A,T,G,C,B) using RepNano Model 2. As anticipated since the cells were only labelled for a small fraction of S phase, the percentage of B was < 0.2% in most reads, with more B incorporated after 60 min than 30 min (Fig. 3c). The reads were mapped on the yeast genome after conversion of the Bs into Ts, and B/T density was represented as a color map as described previously (Fig. 3d). Using all the "pass" and "fail" reads, we obtained a similar genomic coverage for the two samples (21.7X and 19.7X), but a higher replicative coverage after 60 min (1.7X) than 30 min (0.56X), as expected. Although these low coverages did not allow to detect enrichment of B dense fragments at single ARSs, aggregate analysis of all early ARSs (as defined by [25]) revealed a clear enrichment of BrdU-dense tracts (called as described in Methods) 30 min after the release in S phase (Fig. 3e). In contrast no enrichment was observed at aggregate late ARSs. More BrdU-dense fragments were detected 60 min after release (Fig. 3d), and those fragments were more extensively spread around early ARSs (Fig. 3e), consistent with replication fork progression during S phase.

Taken together, our results show that it is possible to precisely map replicative incorporation of BrdU using nanopore sequencing and neural networks trained on appropriate datasets.

## DISCUSSION

BrdU is broadly used to label replicating DNA in yeast and mammalian cells [26, 27]. We observed that growth of MCM869 yeast cells was much slower in the presence of BrdU than thymidine. However, when synchronised cells in G1 were released into S phase in the presence of thymidine or BrdU, their progression through the first S phase was similar, whereas cells released in the absence of thymidine or BrdU were stalled in S phase (Supp. Fig. S2b). Similar results were previously obtained in a comparable strain [28]. We conclude that the presence of BrdU does not initially perturb S phase progression and that short pulses of this analogue are suitable for replication studies. The long-term effect of BrdU on yeast growth may arise from problems in mitosis and/or replication of BrdU-containing parental strands in subsequent S phases, which is not a concern if BrdU incorporation is limited to short pulses as it is the case for classical singlemolecule analysis. Moreover, BrdU can enter mammalian or MCM869 yeast cells without any permeabilisation, which makes experiments far easier and more precise than incorporation of fluorescent dNTPs as for nanochannel imaging [4, 5].

**Figure 3:**
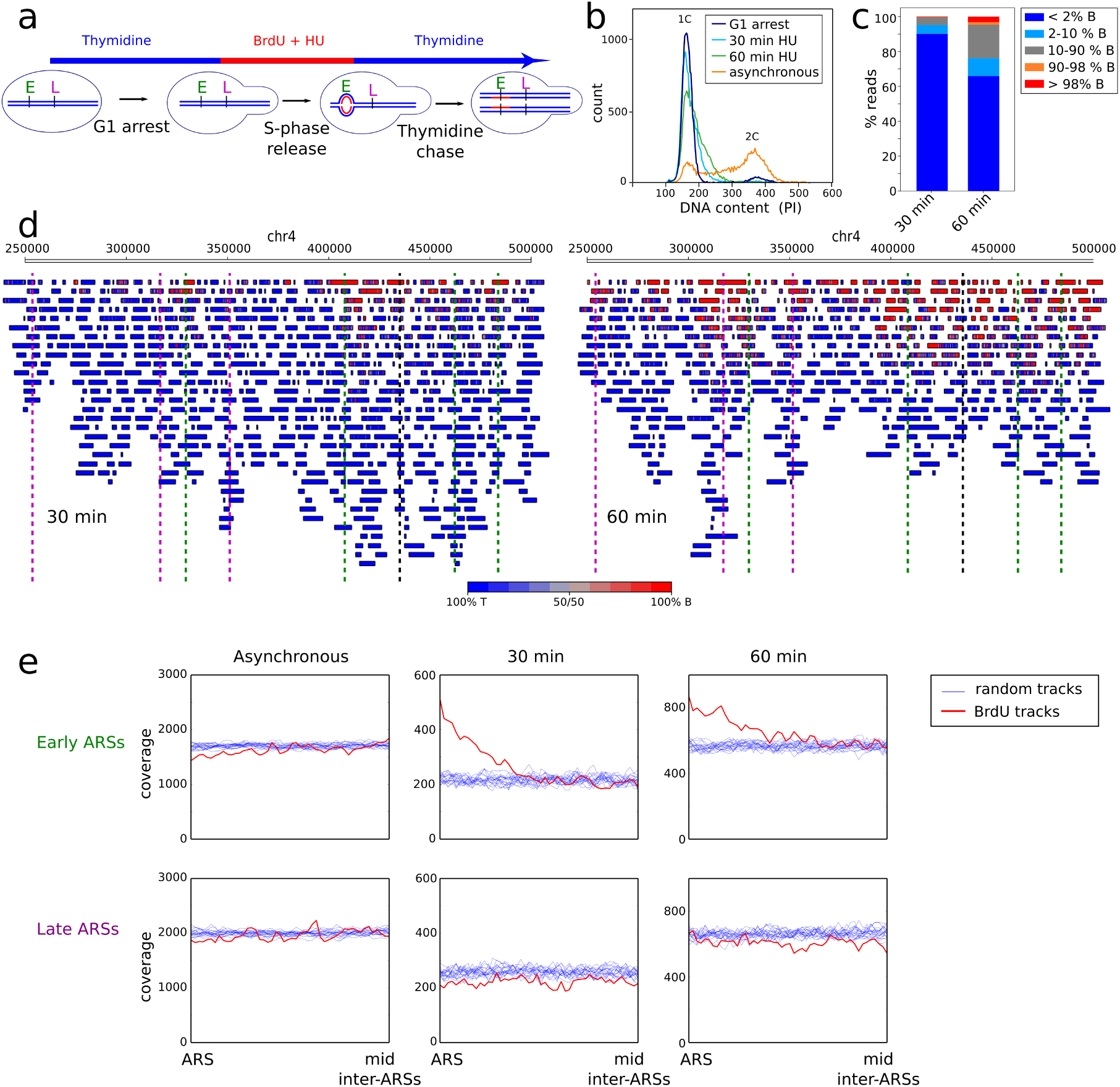
Mapping of early origins in yeast. a. Scheme of the experiment. E, early replication origin; L, late replication origin. b. Cellular DNA was stained with propidium iodide (PI) and cells analysed by flow cytometry to monitor G1 arrest and S phase progression in the indicated samples. c. Proportion of reads with different amounts of B for samples labelled for 30 or 60 min. d. An exemplary 250 kb chromosomal segment with reads from the 30 min (left) or 60 min (right) samples. ARS positions are indicated as dashed lines (green, early; magenta, late; black, unknown timing [25]). e. Coverage of BrdU dense tracts (red) averaged on rescaled inter-ARS regions using either 109 early ARSs (top pannels) or 127 late ARSs (bottom pannels) [25]. ARSs were aligned on the left side of each panel. The coverage of 20 sets of random tracts with the same size distribution is plotted in blue. All "pass" and "fail" reads were used for this figure.

We demonstrate here that the presence of BrdU in the DNA influences the nanopore sequencing signal. The width of current value distribution at specific positions is 5-10 pA. The current shift induced by BrdU substitution for thymidine is within the same range, meaning that B and T current distributions at a single position overlap. As a consequence, Bs or Ts cannot be called with certainty at all positions. This is not problematic inasmuch as our goal is to identify replicative tracts of hundreds of bases but this will limit the resolution of tract ends to ~20-100 bases. This achievement already constitutes a significant improvement compared to optical methods where reliable length measurement of replicative tracts < 3 kb is not feasible.

Good training datasets are critical for successful machine learning. Using a yeast strain dependent on exogenous thymidine or thymidine analogues to replicate its genome, we obtained a broad variety of either native or fully BrdU-substituted DNA fragments. The calling by Metrichor was good enough to map the BrdU-substituted DNA fragments so that pairs of current and reference DNA sequences could be given as training inputs. We implemented RepNano, a neural network basecaller following the architecture proposed in DeepNano [9] to generate a model that was initially trained assuming that every T site had incorporated a B in the MCM869 sample grown in BrdU medium. This first model was good enough to reclassify the reads as substituted or not and to generate a second and more accurate model after reallocating the unsubstituted reads from the BrdU sample to the correct training dataset.

We then tested this model on DNA from cells synchronously released into S phase in the presence of BrdU and HU to selectively label early ARSs. We obtained BrdU tracts that were localised at the expected loci. This result demonstrates the efficient identification of the canonical bases as well as of BrdU by our basecaller.

Human replication origin identification has been controversial due to inconsistencies between techniques and laboratories [29]. In principle, our neural network model trained on MCM869 should be directly applicable to human DNA and allow mapping of early replication origins from synchronized human cells, with nanopore sequencing throughput as the only remaining limitation. With a single MinION run, we covered ~20 times the genome of *S. cerevisiae* and obtained a replicative coverage between 0.5X and 1.7X. This is already an enormous improvement compared to traditional single-molecules techniques such as DNA combing, where months of work are required to collect and analyse a few hundreds of replication events covering a 1Mb locus. Since the experiments presented in this paper were performed, the throughput of a MinION run has increased ~10 fold, and new devices such as the GridION (5 chips in parallel) and the PromethION (48 chips) have been released by ONT. Obtaining a new map of human early origins by nanopore sequencing is therefore within reach.

When mapping replication in non-synchronized cells, it becomes necessary to determine replication direction along labelled tracts so as to discriminate between initiation, termination and elongation signals. To do so, cells are typically consecutively pulsed with 2 different nucleoside analogues. Importantly, in addition to BrdU, several thymidine analogues are available, among which iodo-, chloro- or ethynyl-deoxyuridine have been widely used in DNA replication studies. Future experiments will reveal which analogues alter the nanopore electric signal in a way that allows to discriminate them from thymidine and from each other.

## CONCLUSIONS

This work demonstrates for the first time the power of nanopore sequencing to study DNA replication at the single-molecule level following replicative incorporation of BrdU. This represents an important step forward compared to current single-molecule methods. This paves the way to genome-wide, single molecule replication mapping studies at unprecedented high resolution, which will likely transform this research field.

## MATERIALS AND METHODS

### Primer extension

pTYB21 plasmid (New England Biolabs, NEB) was linearised with EcoRV (NEB) and purified using home-made SPRI (Solid Phase Reversible Immobilization) beads. After initial denaturation for 5 min at 94°C, primer extension was performed on 200 ng of linear plasmid, using 0.5 U/ μL LongAmp Taq DNA polymerase (NEB) with 300 μM of each dNTP (with either dTTP or BrdUTP (Thermo Fisher Scientific, TFS)) and 400 nM of NanoP-pTYB-F primer (5’-ATCGTCGACGG ATCCGAATTCCCTGCAGGTAATTAAATAACTAGTTGATCCGGCTGCTAACAAAGCCCGAAAGGAAGCTGAGTTGGCTGCTGCCACCGCT-3’; Eurofins) for 20 min at 65°C. A control (‘water’ lane in Fig. 1b) with only dATP, dGTP and dCTP was also included. The ssDNA was then digested by ExoSAP iT (TFS) for 30 min at 37°C and the DNA was purified using SPRI beads. The size of the product was assessed using an Agilent DNA 12000 chip on a Bioanalyzer and its amount was quantified using Qubit dsDNA HS Assay (TFS).

### Yeast

MCM869 genotype is *MATa ade2-1 trp1-1 can1-100 leu2-3 his3-11,15 URA3.:GPD-TKyx AuR1c::ADH-hENT1 bar1::LEU2 cdc21::kanMX*. MCM869 was grown in minimum medium (Synthetic Dropout Base, DOB, with Complete Supplement Mixture, CSM; MP Biomedicals) with 100 μM thymidine (Sigma-Aldrich).

To synchronise MCM869 cells in the G1 phase of the cell cycle, cells from an overnight culture were grown for 1h30 in fresh minimum medium with thymidine and synchronised in G1 by incubation for 3h in the presence of 0.2 μM α-factor (Sigma-Aldrich). Synchronisation was confirmed by visual inspection of the cells with a microscope, as MATa yeasts respond to α-factor by growing a projection with a distinctive shape known as a shmoo. To check the impact of BrdU on the first S phase, G1-synchronised cells were washed and incubated in minimum media with α-factor supplemented or not with 100 μM thymidine or BrdU for 30 min before release into S phase by addition of Pronase (50μg/ml, EMD Millipore). Timed aliquots were fixed in 70% ethanol for flow cytometry analyses (see below). For early S phase labelling, the synchronized cells were washed and kept in minimum medium with 0.2 μM α-factor and 0.2M HU supplemented with 100 μM BrdU for 30 min before release into S phase by Pronase digestion. After 30 or 60 min, the cells were washed twice and resuspended in minimum medium with 100 μM thymidine for 1h to convert replication intermediates into dsDNA. The cells were then pelleted and frozen. DNA was purified by Zymolyase, RNAse A and proteinase K digestion followed by phenol-chloroform extraction (samples from Fig. 2) or using Qiagen Genomic-tips according to the manufacturer instructions (samples from Fig. 3). This second method gave longer read sizes (Supp. Fig. S5a). The size of the DNA was checked by agarose gel electrophoresis and on an Agilent TapeStation.

### Flow cytometry analysis

Cells in 70% ethanol were washed with PBS, treated with 0.25 μg/μL RNAse A for 1h at 50°C, and with 1.8 μg/μL proteinase K for an extra hour at 50 °C. Cells were stained with 100 μg/ml propidium iodide (PI) in PBS for 15 min in the dark, resuspended in 5 μg/ml PI, sonicated and analysed using a Cytoflex flow cytometer (Beckman). 20,000 events were recorded for each sample, the doublets and the debris were filtered out for cell cycle analyses.

### Nanopore sequencing and data processing

MinION sequencing libraries were prepared according to the manufacturer protocols. For the plasmid, the 2D low input kit with R9 chemistry was used. For the yeast DNA, the 2D kit with R9.4 chemistry was used. Details about ONT protocol and software versions used for the different samples can be found in Table S1 and S3. Raw reads were basecalled using Metrichor (ONT) with default parameters and aligned on the reference (plasmid or S288C genome) using BWA MEM [13] with the -x ont2d option (-k14 -W20 -r10 -A1 -B1 -O1 -E1 -L0). In order to visualise current values at a specific genomic loci, the event sequences were extracted using poretools [30] and aligned on the reference using the sam file information and custom Python scripts. Current values were normalized subtracting the whole read average current. The PCA plot was generated using R pcaMethods library.

### RepNano

We implemented RepNano, a neural network approach with the same architecture as the one proposed in DeepNano [9] to convert the raw current from nanopore experiments in a sequence of bases. As in DeepNano, the input of RepNano neural network is the output from Metrichor, which segments the raw nanopore current in plateaus of different lengths (typically from 5 to 10 current values). Each plateau P_i_ is characterised by its mean value m_i_, its standard deviation s_i_ and its length l_i_. In fact, m_i_ and s_i_ are rescaled and the input of the neural network is a 4-vector containing the rescaled mean, the square of the rescaled mean, the rescaled standard deviation and the length of the plateau 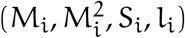. The rescaling is performed on the whole sequence input as follows:

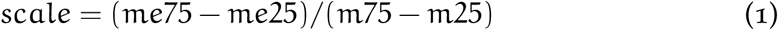

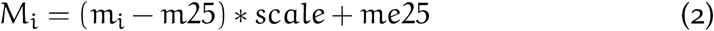

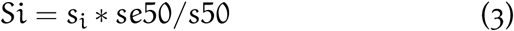
with m25 = percentile(m_i_,25) and m75 = percentile(m_i_, 75), s50 = median(s_i_), me25 = 0.07499809, me75 = 0.26622871 and se50 = 0.6103758.

The neural network is composed of three bidirectional Recurrent Neural Network (RNN) and one time-distributed softmax layer (Supp. Fig. S3a). Each bidi-rectionnal layer is composed of two long short-term memory (LSTM) [31] layers with n = 20 hidden units (100 units in DeepNano), that respectively process the input in forward (F^(n)^) and backward (B^(n)^) directions, and then concatenate both outputs (O^(2n)^) (Supp. Fig. S3b). The number of parameters for each layer is detailed in Supp. Fig. S3c. The final softmax layer outputs a vector of length 6, normalised to 1, that represents the probability to have A,T,C,G,BrdU(B) or no base. Here the architecture is simplified with respect to DeepNano that outputs two vectors, allowing a plateau to be eventually associated to two bases. RepNano was implemented in Python with the Keras library [32] and is available at https://github.com/***.

### RepNano training

In DeepNano, the loss function used for the training requires a one-to-one alignment between the sequence of current plateaus and the DNA sequence in the training sets. To reduce the impact of imperfect mapping, RepNano uses a connectionist temporal classification loss function [33] that does not require this one-to-one alignment. This loss function was developed in the context of speech recognition and was designed to map a spectrogram sequence to a word sequence, even with a non-perfect alignment between them. The DNA sequence used for the training is the sequence predicted by Metrichor, corrected after alignment to the yeast reference genome. We used two datasets for the training: a thymidine dataset with normal DNA where all the sequences have only A,T,C,G bases, and a BrdU dataset where all Ts are replaced by Bs in the target sequence. The training was done on sequences of 4-vectors of length 40 (series of 40 plateaus) and the error between the prediction and the correct sequence was minimised using a stochastic gradient descent optimizer implemented in Keras [32]. 1000 iterations were used for each model.

### BrdU tract calling and coverage plots

After basecalling with the different models, the reads were mapped as above after conversion of Bs into Ts. Files containing all the reads with the B and T positions on the genome were generated using sam file information and a custom Python script and were used to calculate and plot B density along the reads as a colormap for visualisation. BrdU dense regions were called using a running mean smoothing (w = 50 B/T) and two thresholds : (i) the maximum density of a called tract should be > 0.8 (80% of B), (ii) the borders of the called tract should be > 0.6. The coverage of these called tracts relative to the ARSs was calculated by summing up all the inter ARS regions using either early or late ARSs [25]. Control sets of random regions with the same size distribution were also plotted. The rDNA region (chr12 : 440000-480000) was excluded from this analysis as these repeats strongly affected the averaged signal.

## DECLARATIONS

### Availability of data and material

The datasets generated during the current study are available from the ENA repository [PRJEB28657]. RepNano can be downloaded from https://github.com/***.

### Author’s contributions

OH designed the project and raised fundings. MH, FP and BLT performed the experiments. CC, SE and AL performed the nanopore sequencing. JMA and BA implemented RepNano. MH and EN analysed the data. MH and OH wrote the manuscript with inputs from the other authors.

## Acknowledgements

The authors thank all members of O.H. lab as well as Alain Arneodo, Patrice Abry and Nelly Pustelnik for helpful discussions.

